# Folate bioavailability in reconstructed skin models and effects of folate in a monolayer wound healing assay. Approaches on topic application

**DOI:** 10.1101/2021.04.07.438798

**Authors:** Dirk Dressler, Martin Ulmann, Gerd Wiesler

## Abstract

In chronic and degenerative diseases impacting the skin folates play an important metabolic role improving wound healing and reducing skin irritations. In contrast to systemic folate administration little is known on skin penetration of folates after topical application. Here the penetration of simple aqueous solutions of reduced folates have been investigated with in-vitro reconstructed skin models mimicking the barrier of native human skin. For up to 24 h, penetration of the epidermis by newly developed folate salts and formulations were investigated. Aqueous and lipophilic solutions of L-formyltetrahydrofolate and L-mefolinate salts were able to penetrate the epidermis. Even more importantly, the skin model revealed the metabolic conversion of L-folinate to L-methyltetrahydrofolate. Exemplarily the effects of these new folate salts have been tested on wound healing in a scratch assay with primary human keratinocytes. All folates applied were able to enhance wound healing compared to the control.

Bioavailability, metabolic conversion and physiologic effectiveness of new folate formulation have been shown successfully in in vitro applications providing evidence for the potent applicability of the new folates in preparations for topic application.

## Introduction

The skin much more than other tissue of the body is exposed to detrimental environmental conditions. Sunlight, smoke or air pollution [1], microorganisms, and chemical challenges like household detergents amongst other influences challenge the health of the skin [2]. Besides the extrinsic factors the condition of the largest organ is affected by pure intrinsic aging regularly, enhanced by photo aging processes not only associated with reduced number of blood vessels, especially in the upper dermis [3]. Consequently, regenerative abilities of our first barrier skin increasingly are affected with age [3–5]. Additionally, wounds caused by various incidents more or less traumatically inflict the barrier between the body and the surrounding. Due to the initial great regenerative potential of the skin a lot of the integrity losses are cured by proliferation and differentiation of the cellular elements of the skin [6, 7]. Wound healing can be divided into several phases such as inflammation, proliferation, and remodeling and age-related changes in wound repair have been described in each of these phases [8–10]. The underlying mechanisms remain unclear. Skin keratinocytes from older donors have more limited replicative lifespan than keratinocytes obtained from younger individuals when placed in culture [11, 12]. Cell adhesion molecules of estrogen levels were identified as possible factors influencing impaired wound healing in elderly people [13–15]. Not exclusively but with increasing importance in the aged there are also metabolic dietary factors that may affect skin conditions. Among the most important nutrients for human health the family of folates play a key role in human skin. “This importance is underscored by potential links between folate deficiency and psoriasis, vitiligo, exfoliative dermatitis, glossitis, and skin cancers.” [16].

Besides the metabolic benefits of orally taken folates, it would be interesting to understand if folates can cross the skin and thus open up the path for an alternative route of entry simultaneously offering the possibility to apply folates directly to areas that have increased needs.

The main drawbacks associated to folic acid use, particularly for topical applications, are the limited solubility and the sensitivity to UV rays [17]. The very low solubility of folic acid in aqueous physiological solutions (1.6 mg/L) [18] results in insufficient homogeneous dispersion in hydrophilic solvents. Therefore, surfactants, co-surfactants or co-solvents are required to employ a homogeneous formulation. The insolubility in organic solvents and the low lipophilicity hinders the penetration of the skin while applying penetration enhancers may impair the skin barrier.

Folic acid is known as a food supplement for a long time. Folic acid is a synthetic form of the vitamin, which is only found in fortified foods, supplements and pharmaceuticals. It lacks coenzyme activity and must be reduced to the metabolically active tetrahydrofolate form within the cell [19].

Newly developed reduced folate-salts or folate formulation with improved solubility characteristics have been considered to investigate their ability to penetrate the skin in aqueous solution. These reduced folate salts (L-Formyltetrahydrofolate di arginine abbr. L-FTHF di arginine, L-Methyltetrahydrofolate di choline abbr. L-MTHF di choline) are more lipophilic, show good stability, are readily soluble in aqueous solutions, and are at least moderately soluble in organic media increasing the options for formulation for local applications. The solubility characteristics of L-Formyltetrahydrofolate Calcium (L-FTHF Ca, approx. 0.51 g/L in aqueous solutions) has been improved significantly in a new proprietary liquid formulation avoiding precipitation of the folate.

The basic question motivating the current approach was the bioavailability of the new folate salts and possible differences in uptake efficiency between the different formulations. Based on these results the development of products for topical application might be considered. To address at least one of the fields of application for such products the effect of these specific folate salts on wound healing was evaluated on behalf of the scratch assay including primary skin keratinocytes.

## Material and Methods

### Chemicals

L-FTHF Ca (Fig 1; L-formylfolate calcium salt, 5-formyl-(6S)-tetrahydrofolic acid calcium salt) purchased from Cerbios SA, Switzerland; L-MTHF di choline (Fig 2; L-methylfolate di choline salt, 5-methyl-(6S)-tetrahydrofolic acid di choline salt) and L-FTHF di arginine (L-formylfolate di arginine salt, 5-formyl-(6S)-tetrahydrofolic acid di L-arginine salt were synthesized by SynphaBase AG, Switzerland on behalf of Aprofol AG, Switzerland.

**Fig. 1:**
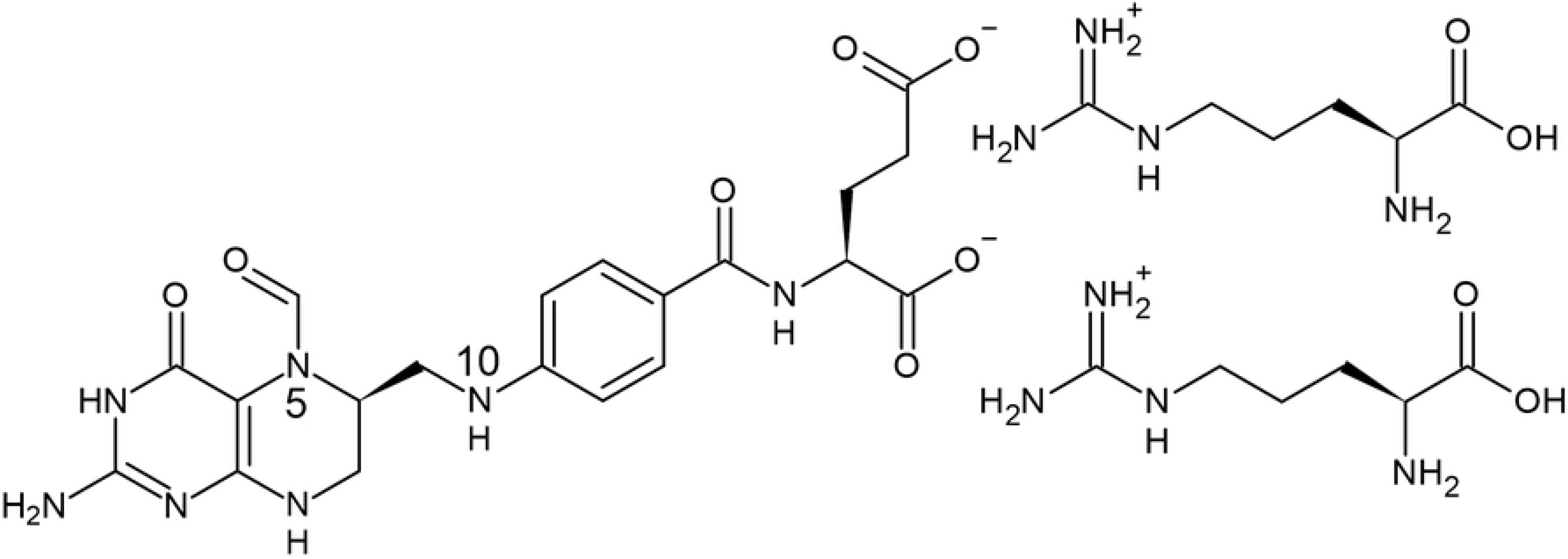
Structure of L-FTHF di arginine

**Fig 2:**
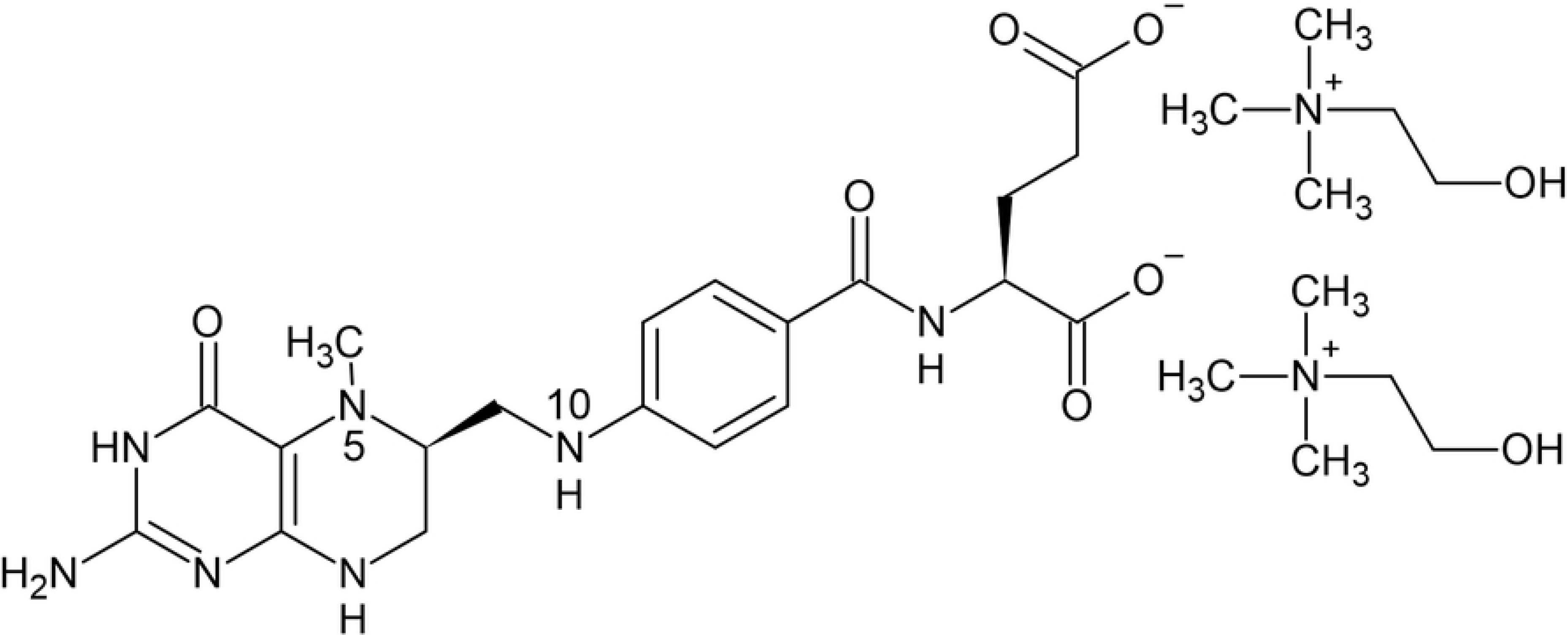
Structure of L-MTHF di choline

Solutions for the assays were prepared at room temperature without pH modification as follows: L-FTHF Ca 2.5% (w/w) and sodium gluconate 2.5% (w/w) in ultra pure water (MilliQ Reference A+, Merck, Darmstadt, Germany) corresponding to 48.69 mM L-FTHF. L-MTHF di choline 2.5% (w/w) in ultra-pure water corresponding to 37.55 mM L-MTHF. L-FTHF di arginine 2.5% (w/w) in ultra-pure water corresponding to 30.42 mM L-FTHF.

### Bioavailability

Bioavailability of folates in skin, was evaluated on behalf of reconstructed skin models (epiCS, Henkel AG & Co. KGaA, Düsseldorf, Germany) generated from isolated primary human keratinocytes. These models show a barrier very similar to human skin and are an accepted model for skin irritation and skin corrosion tests (e.g. OECD TG 439).

The skin models are cultured on a porous membrane forming a two-compartment system separated by the human skin like barrier (0.6 cm^2^). The skin models are delivered accompanied with quality control data of the respective batch. Additionally, to determine the applicability of the skin models at the study site, the barrier was checked for leaks before use by means of trans epithelial electrical resistance measurements (TEER, EVOM with STX3 electrodes, World Precision Instruments, USA). Models with a significantly below-average electrical resistance between the apical and the basal compartment were not used in the experiment. Suitable skin models were treated topically with 50 μL of the test product solutions prepared as described under “chemicals” without further dilution. The basal compartment during treatment was filled with 1000μL HBSS (Hanks balanced salt solution Lonza, Switzerland) to omit influences of the complex and proprietary cell culture medium on later analyses. After 4h (data not shown), 8h and 24h, samples of the buffer (500 μL of 1000 μL) were taken from the basal part below the models to check for the presence and concentration of folates. 500 μL of fresh HBSS was added replacing the withdrawn volume at 4h and 8h. Each test product and controls were tested at one concentration in three replicates. The study was repeated once to obtain results from two independent trials.

The analysis of the buffer samples was carried out by way of a LC/MS method established by the University of Saarland, Germany [20].

Subsequently to the topical treatment for 24h the skin models were washed and incubated with a vital dye (Resazurin, MerckMillipore, Darmstadt, Germany) to determine the relative number of living cells in the models. After 2h incubation samples of the basal medium were analyzed on Resarufin the redox product of Resazurin formed by vital cells by measuring the fluorescence at 560/590 nm on a fluorescence reader (Infinite M200 pro Tecan, Austria).

### Scratch Assay

To exemplarily examine the effect of different reduced folate preparations on wound healing the in vitro Scratch Assay was employed. This in vitro model comprises the defined injuring of a closed cell layer and the subsequent recovery of the defect in the presence of the test-substance compared to appropriate controls, i.e. (untreated) cells with no test-substance present.

The experimental procedure for the Scratch Assay was as follows: Primary keratinocytes of human skin (C-12005, Promocell, Heidelberg, Germany) were cultured in the medium recommended by the manufacturer (C-20011, Promocell, Heidelberg, Germany). A dose finding experiment was conducted prior to the scratch assay. Keratinocytes were seeded on 96-well plates cultured to confluence and incubated with five dilutions of the folate solutions for 24 h. MTT (methylthiazolyldiphenyl-tetrazoliumbromid, 0.5 mg/mL M2128, Merck, Darmstadt) was added for 2h and the color change caused by the conversion of the vital dye in living cells was quantified photometrically.

Three different non-toxic dosages were applied in the scratch assay. Keratinocytes were cultivated in appropriate cell culture plates (6-well plates, Greiner BioOne, Frickenhausen, Germany) bearing position markings on the outer bottom of each well. Cells were grown until they completely cover the growth area. Cells were scratched with a pipette tip along the position markings, generating an area free of cells (scratch, injury). The cultures were washed to remove partially detached cells and were covered with medium comprising the test-substances in appropriate dosages. Regrowth of the cells into the cell free area was photographically documented at 0h, 6h, and 24h. Analysis was done by marking and calculating the cell-free area on behalf of a software (ImageJ [21]) on the image files taken at the given points in time. The relative closure of the wound-area was calculated by subtracting the relative wound area at each time point from the initial wound area (=100%). For every dosage of each test substance two replicates were treated in a single experiment. The not supplemented standard medium and standard medium supplemented with 10 % fetal bovine serum (FBS, Biochrome, Berlin, Germany) served as control.

## Results

The bioavailability of L-MTHF or L-FTHF from the preparations was demonstrated after supplementation of reconstructed skin models. Buffer samples of the basolateral compartment of the models were analyzed for folic acid, L-MTHF (L-methylfolate), and L-FTHF (L-formylfolate) in order to detect the penetration and metabolism of the different folate salts.

After 8 h low folic acid concentrations were found partially differing significantly between control models and supplemented models. L-MTHF and L-FTHF were found in much higher concentrations. After supplementation, the topically applied folate derivatives penetrated through the epithelium of the skin models and partially were metabolized by the cells of the skin models. The main part of the topically applied L-FTHF passed through the tissue but was also partially metabolized to L-MTHF (Fig 3, and 4). Conversely, the topically applied L-MTHF as such got through the tissue of the skin models obviously without being metabolized.

**Fig 3:**
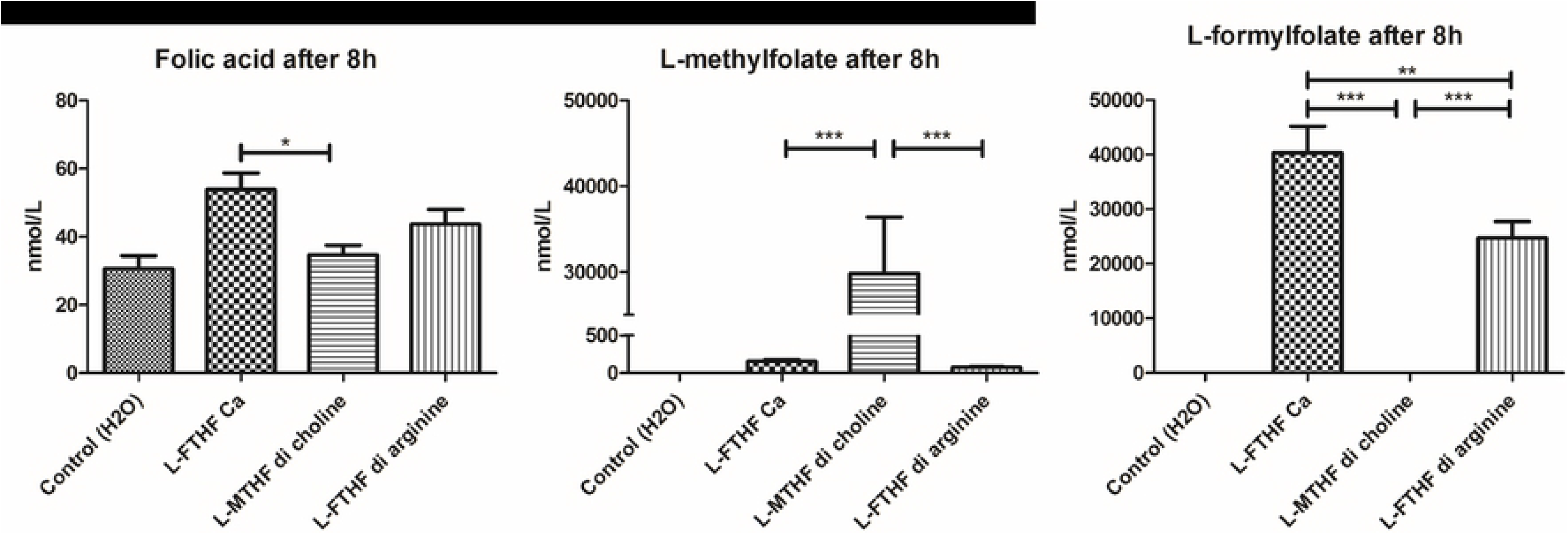
Penetration of folates through reconstructed human skin models. Determination of folic acid, L-MTHF and L-FTHF concentrations in the compartments under the skin models after 8h. Skin models were topically treated with the compounds indicated. Mean of two experiments each with N=3 + SEM. Statistics: ONE-way ANOVA with Tukeys multicomparison test (Prism 5.04 GraphPad Software, San Diego, USA) (p < 0.05 = *; p < 0.01 ‘= **; p < 0.001 = ***). Differences to the control are only partially depicted.

**Fig 4:**
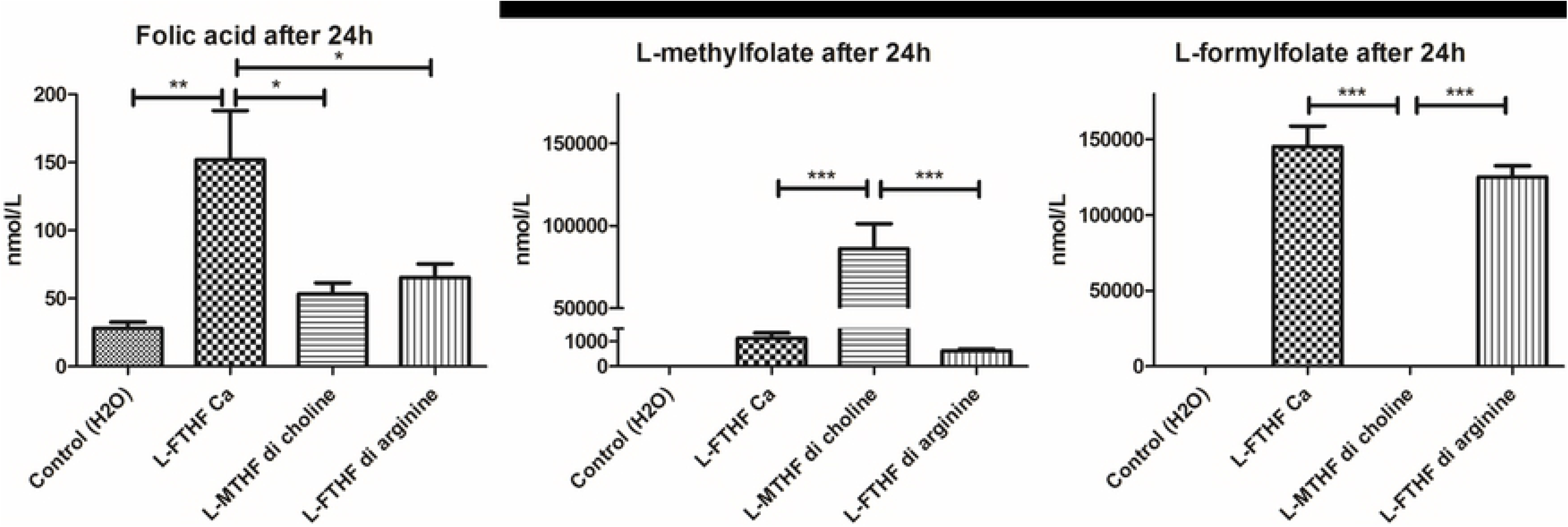
Penetration of folates through reconstructed human skin models (24h) Determination of folic acid L-MTHF and L-FTHF concentrations in the compartment under the skin models after 24h. Skin models were topically treated with the compounds mentioned on the x-axis. N=3 + SEM. Statistics: ONE-way ANOVA with Tukeys multicomparison test (Prism 5.04 GraphPad Software, San Diego, USA) (p < 0.05 = *; p < 0.01 ‘= **; p < 0.001 = ***)Differences to the control are only partially depicted.

After 24h (Fig 4) both the folic acid (partially) and L-MTHF concentrations are higher than after 8h. The folic acid concentrations below the skin models treated with L-FTHF Ca are twice as high as the concentrations of the other skin models. However, the folic acid concentration is negligible in comparison to L-MTHF and L-FTHF concentrations measured (refer to the molar range). The differences in the level of L-MTHF between L-FTHF Ca and L-FTHF di arginine supplemented on the one hand and L-MTHF di choline supplemented skin models were highly significant (Figs 3 and 4). The concentration of L-MTHF in skin models treated with L-FTHF Ca also are relatively high (1118 nmol/L). The concentration of the solution below the models treated with L-FTHF di arginine is approximately half that (620 nmol/L). L-FTHF cannot be detected in the samples after L-MTHF di choline since L-FTHF cannot be formed from L-MTHF by human cells or only through energy-consuming metabolic pathways (Figs 3 and 4). On the other hand, L-FTHF is measurable in all other samples with the exception of the control.

Since the integrity of the skin like barrier is important to prevent leakage of topically applied supplements to the basolateral acceptance compartment, we examined the vitality of the skin models after each experiment besides the electrical resistance controlled prior to the application of the supplements (data not shown). The viability of the skin models was not adversely affected by the treatment (Fig 5). A slight decrease in vitality following treatment with the L-MTHF di choline was observed (86% of the control). However, a decline of this magnitude is not considered being a toxic effect.

**Fig 5:**
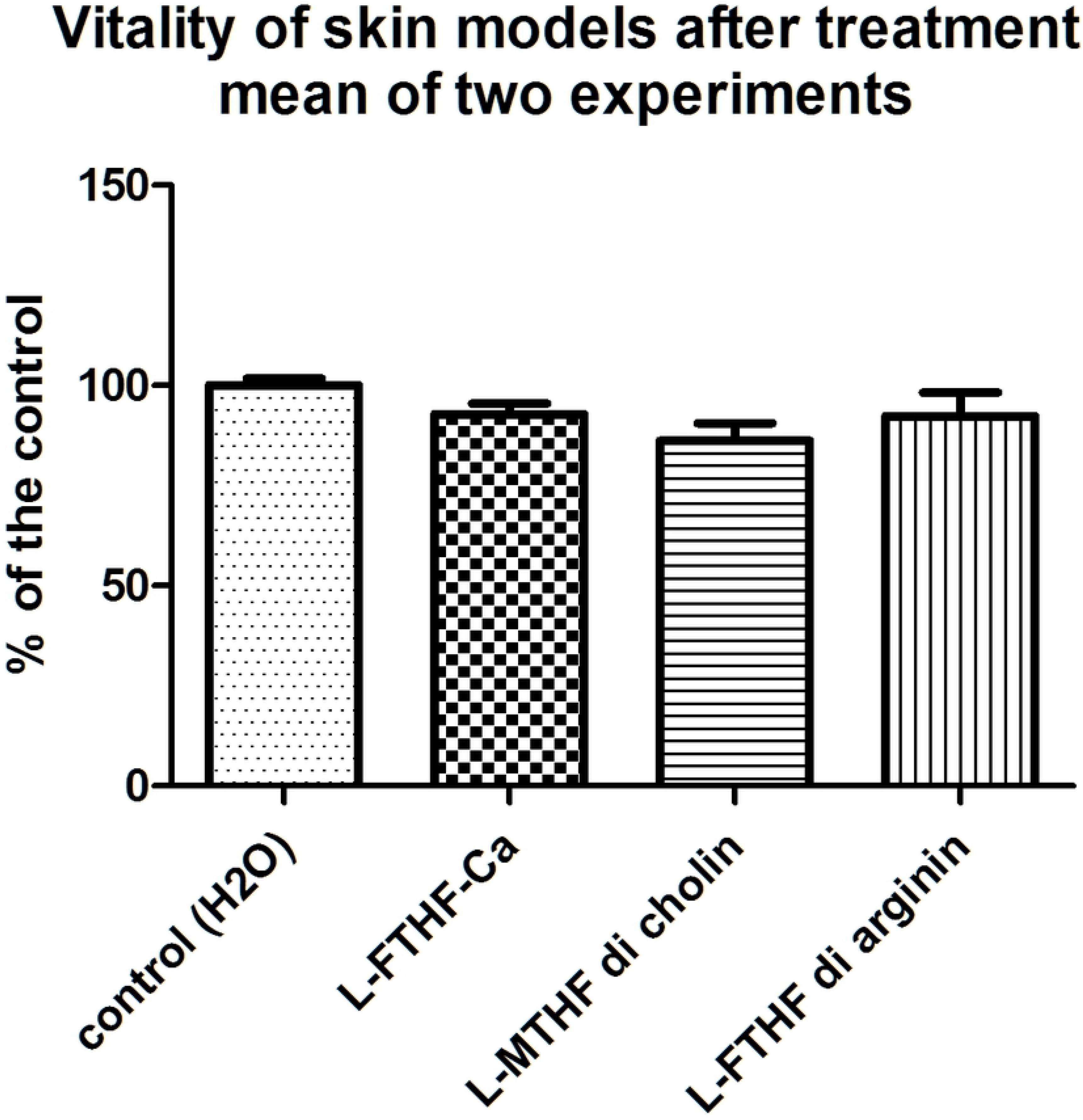
Survival of reconstructed human skin models. Relative vitality of the cells of the skin models after 24h topical treatment with 50 μL of each of the test substances or water. The fluorescence values of the dye (resazurin) is corresponding to the number of living cells and was normalized to the control (= 100%). Mean of two experiments each with N = 3 + SEM.

In preparation to the scratch assay, applicable concentrations were evaluated by applying five different concentrations of the test chemicals to confluent monolayers of primary keratinocytes as described under methods. Concentrations resulting in relative vitalities of more than 70% compared to the control qualify for the scratch assay.

The results for the dose finding are depicted in Fig 6. Based on these results the highest of the three concentrations chosen for the scratch assay were 1.5 mM for L-MTHF di choline, 1.95 mM for L-FTHF Ca and 1.22 mM for L-FTHF di arginine. These concentrations were supported by additional calculations (Hill-slope, GraphPad Prism 5.04, data not shown).

**Fig 6:**
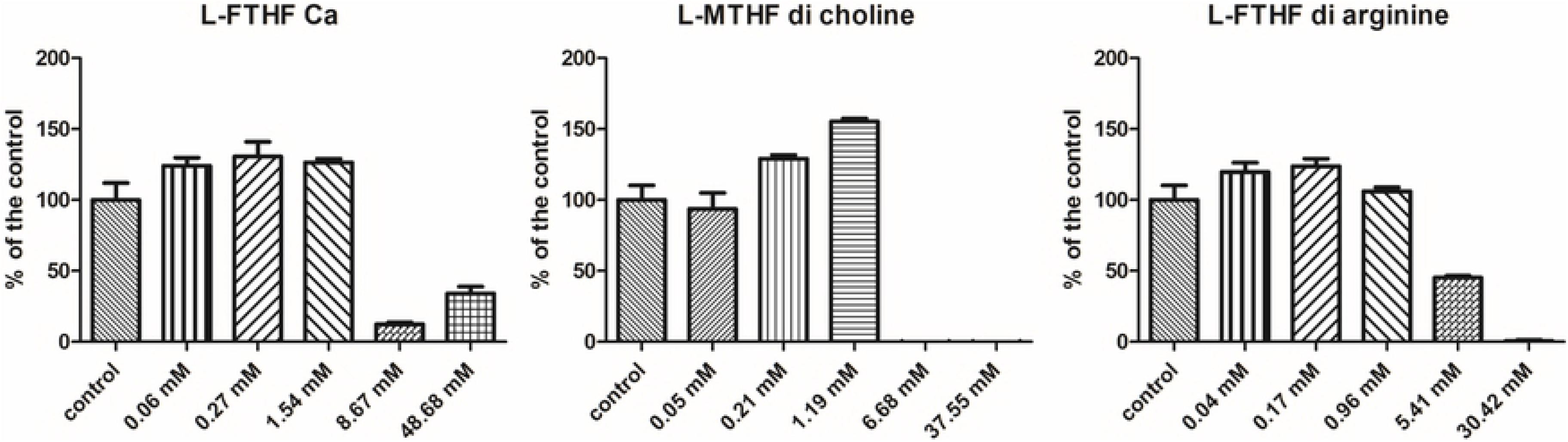
Determination of the applicable dosage of the test products. Cytotoxicity test with keratinocytes in monolayer culture after conversion of the vital dye MTT by living cells. The concentration data are based on solutions as mentioned in chemicals. Treatment time = 24h. Untreated cultures served as negative control. N = 6 + SEM.

Following the procedure described under methods, onto confluent monolayers of primary keratinocytes, a wound was applied and the three different concentrations of the test chemicals were added to separate cultures. Other keratinocyte cultures were treated with medium only or medium with 10% FBS accordingly and served as controls.

After 6 hours, no substantial coverage of the cell-free area has been achieved in all treatment groups. Only 12 to 24% of the initial area were recovered. After 24 h, 50 to 77% of the initial wound area was covered. At that time the folic acid solutions in tendency better promote growth than the standard medium (Figs 7). The folate solutions promote growth in a similar manner (60 - 77% recovery). No clear dose dependency was detected with L-FTHF di arginine. No statistics have been calculated due to the low number of replicates.

**Fig 7:**
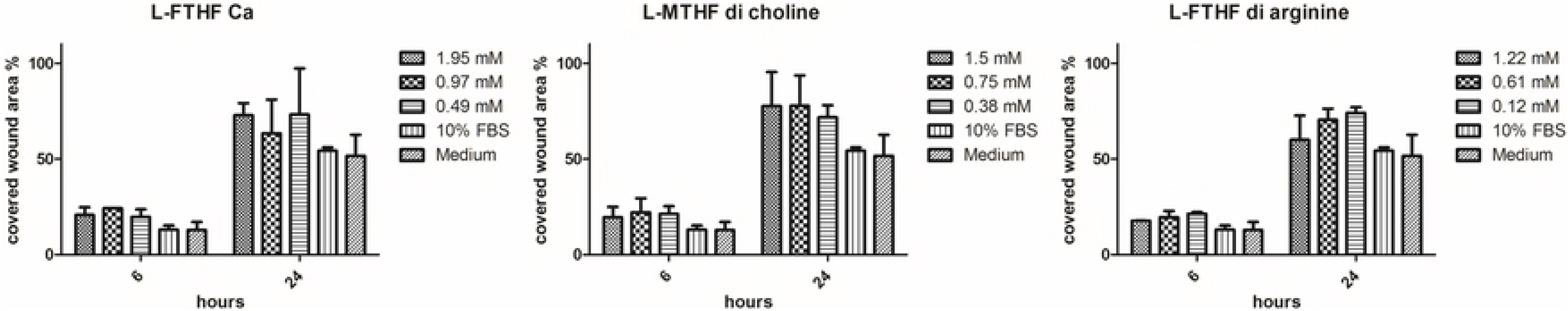
Scratch assay, recovery of wound area. Area of the wounds covered after 6h or 24h. The measured areas were expressed as % of the starting area of the respective treatment group. The higher the bars, the more the wound “healed”. For each treatment group, three concentrations were investigated in two separate cultures (N=2) +SEM.

Supplementation with various folate preparations in tendency led to accelerated regeneration of an artificial wound after 24h in the scratch assay.

**Fig 8–10:**
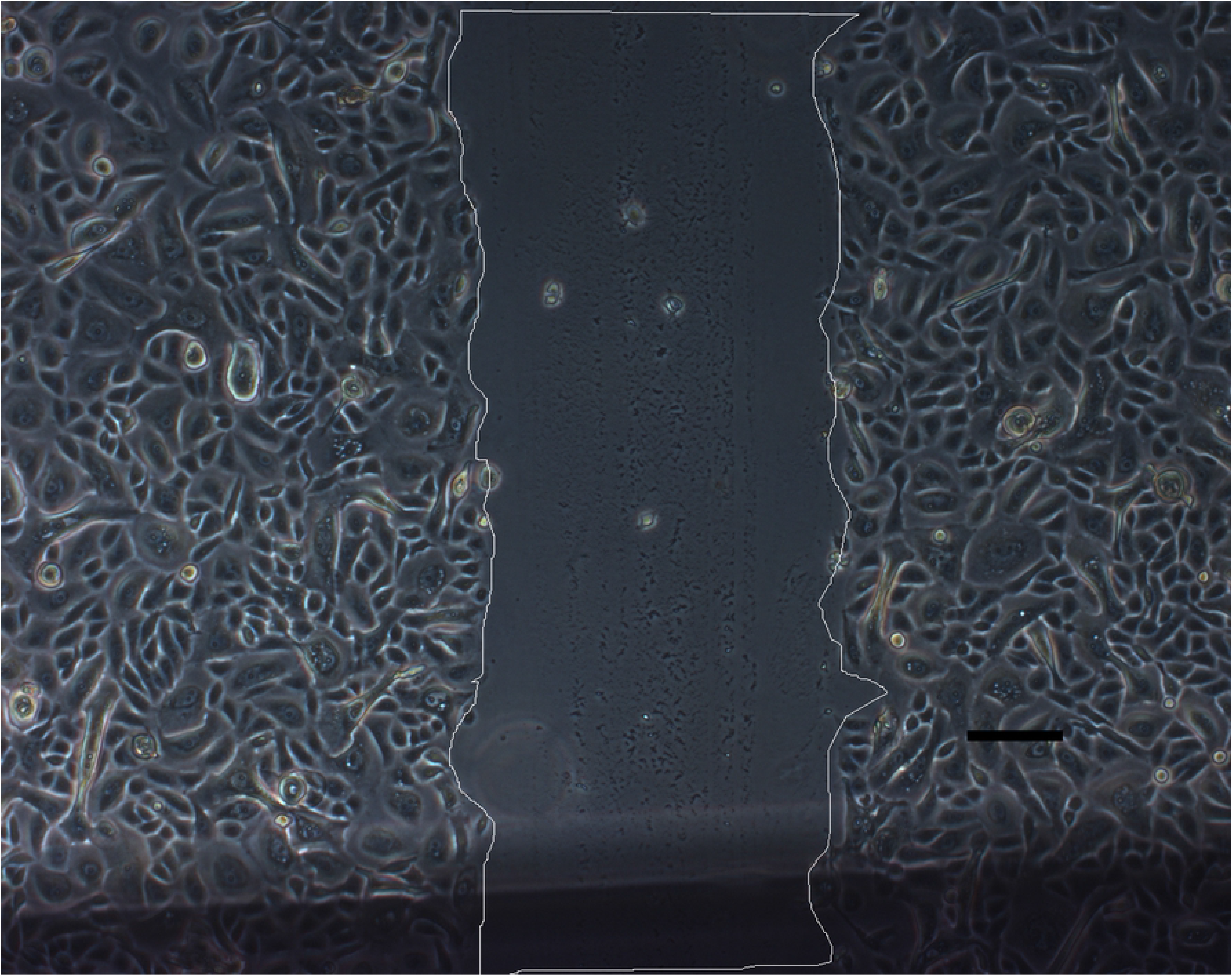
Scratch assay, examples of progression of wound healing. Example of the recordings for L-MTHF di choline 48.9 nmol/mL from left to right 0h, 6h and 24h. The white lines delimit the wound areas for measurement. The black bars mark 100 μm stretch.

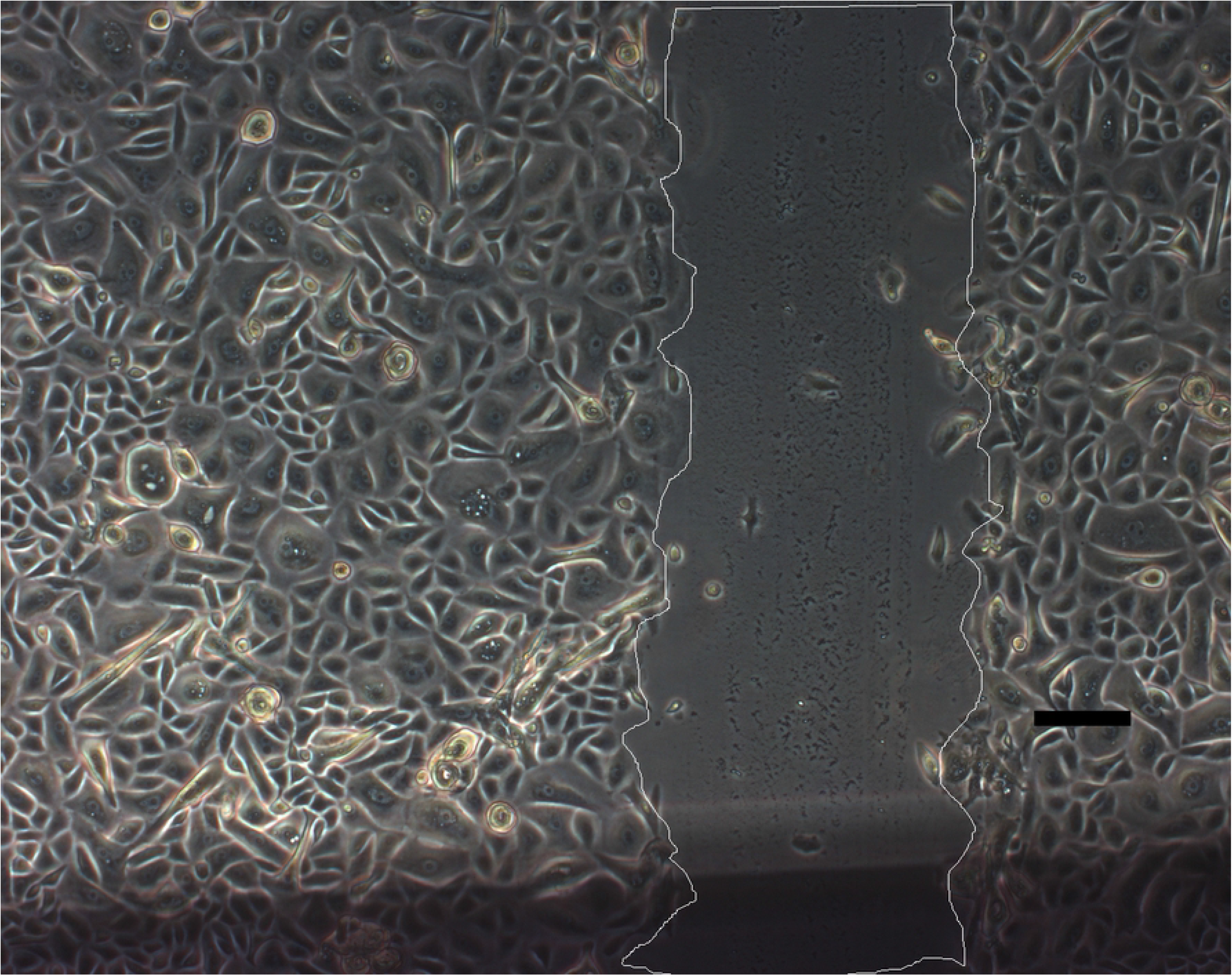

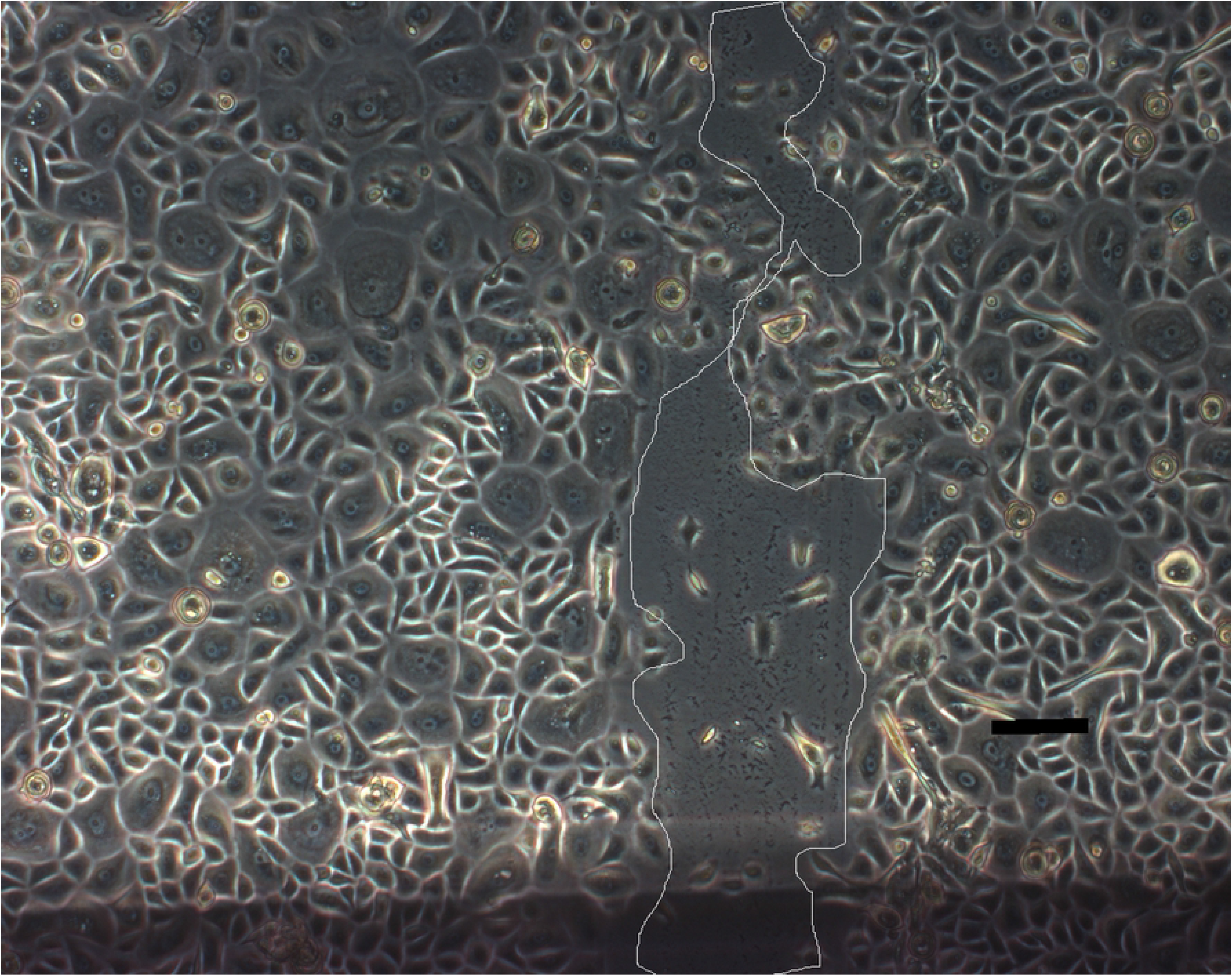

After 6h only little surface recovered. After 24h it becomes difficult to mark a cell-free area. The initial wound area is almost covered.

## Discussion

Hasoun et al [22] found low folate concentrations in human epidermis compared with many other tissues. At the same time they identified a relatively high proportion of L-MTHF in healthy epidermis compared to the dermis. They discuss a special role for L-MTHF in the epidermis with respect to possible photo degradation and maintenance of the high proliferation rate typical for the epidermis. In general these findings indicate special requirements of the epidermis regarding folates.

In the current in vitro study, the bioavailability of newly developed folate salts was demonstrated in reconstructed skin models. This is the first time that penetration of skin models has been shown with simple aqueous solutions of folates free of any allergenic or irritating substances. Most impressing, the supplemented L-FTHF was partially metabolized to L-MTHF the dominant form in the body and the cellular part of the skin. The occurrence of metabolites of the supplemented salts on the one hand underline the accessibility of the folates for human skin cells despite distinct horny layer. On the other hand those findings highlight the physiologic capacity of in vitro reconstructed skin models which largely rebuild the barrier function of human skin.

At both times examined we found similar but low concentrations of folic acid opposite to the treatment side in the basal compartment (Figs 3 and 4). With no other source of folates present than the supplements. We assumed that folic acid values determined with the tissues mainly still originated from folic acid residues fed with the culture medium during maturation of the tissue and now gradually released or leaking from the cells. This interpretation is quite likely for the control tissue that received no supplement. However, the values measured with tissues supplemented with L-FTHF-Ca were significantly different to values measured with the control tissue or tissues supplemented with L-MTHF di choline strongly indicating a possible additional trigger by the treatment. If this finding is caused by lower consumption of folic acid remnants due to a now available second source of folates remains unclear.

L-MTHF was found in high levels opposite from the supplemented side of the skin models. The highest concentrations understandably were found after supplementation with L-MTHF di choline that seemed to penetrate easily through the keratinized layer and the cell layers below. Remarkably, models supplemented with L-FTHF Ca (155 nmol/L after 8h, 1118 nmol/L after 24h) and L-FTHF di arginine (76 nmol/L after 8h, 620 nmol/L after 24h) also showed quite high concentrations of L-MTHF but significantly lower concentrations than after L-MTHF di choline treatment. Obviously, L-MTHF was formed from these two derivatives by cellular metabolism. After 8h and even more after 24h the concentrations of L-MTHF of the L-FTHF-Ca treated skin were twice as high as after L-FTHF di arginine treatment. However, it must be taken into account that molarities applied of the two supplements differ by about one third (L-FTHF-Ca = 48.69 mM; L-FTHF di arginine = 30.42 mM) which contributes to a large part of the difference.The analysis of L-FTHF shows a more simple picture. Since L-MTHF supplemented via L-MTHF di choline can not be metabolized to L-FTHF only L-MTHF has been detected below these skin models. Significantly more L-FTHF was found after supplementation with L-FTHF-Ca than after supplementation with L-FTHF di arginine. As mentioned above this difference to a large part might be caused by different molarities supplemented with these two solutions (see above). After 24h the difference between the supplementation groups were below significant levels. Moreover, the difference of the initially applied molarities no longer are reflected. While after 8h the penetration of L-FTHF-Ca seems to be slightly better, after 24h treatment the contrary becomes evident.

The viability of the models was not significantly affected by the treatment. It therefore could be assumed that the barrier function of the skin models was not impaired. Compared to the test on irritating effects according to OECD TG 439 [23], where models are treated with the test substances for only 15 minutes, treatment over 24 hours is far more challenging. Thus, for the tested folates a very good compatibility to the human skin can be stated.

In the current approach, penetration of folates through the epidermis has been addressed and metabolic conversion was detected as a side effect. Since the reconstructed skin tissue models as such were not examined on folate content it is unknown if and to what extend the folates are retained in the reconstructed skin tissue. Moreover, it is not clear which specific cell layer of the model is responsible for the metabolic conversion of folates. Future approaches might engage more into the processes within the tissue even integrating dermal elements making use of full thickness skin models.

Humans are not able to synthesize folate de novo, and therefore are dependent upon dietary sources. The terms folate and vitamin B9 refer to a large family of chemically similar compounds different in the glutamine residue, one-carbon substituent position at N5 and N10 or the oxidation state. The most widely known folate is the synthetic folic acid due to its enhanced chemical stability [24]. Folic acid is inactive in the body and needs to be converted to reduced folate by several enzymes e.g. in a rate limiting step by dihydrofolate reductase [19, 25, 26]. In further reducing steps dihydrofolate and dietary folates, as monoglutamates, are converted to the bioactive folate form in the body, e.g. L-MTHF. The folate uptake and metabolism is depending on many genetic polymorphisms affecting the status of folate and vitamin B12 resulting in elevated homocysteine [27, 28]. While folate deficiency has been extensively documented by analysis of human plasma, folate status within skin has not been widely investigated. However, inefficient delivery of micronutrients to skin and photolysis of folates argue that folate deficiencies will be present if not exacerbated in skin [24]. Therefore, a targeted delivery of micronutrients might be of future interest to circumvent or support systemic impairments. The characteristics of the selected reduced folates as determined here supports their applicability with this respect.

In addition to the uptake studies aqueous solutions of the selected reduced folates exemplarily showed positive tendencies on wound healing in the scratch assay. This indication of an effect of the tested compounds in wound healing provides hints on a possible field of application improving the integrity of deficient epidermal barrier [22].

Since metabolization of folates as reported for the reconstructed skin models was not examined with the cells in the more basic monolayer model applied for wound healing there still is a lack of information on specific metabolites involved in the observed effects. Whether or not non-differentiated skin keratinocytes in monolayer culture are able to transform folates in a similar manner as highly differentiated keratinocytes in skin models has to be addressed in future work. Results obtained from such trials might help to understand which folates are best choice to promote wound healing or skin related diseases.

## Conclusion

In the skin model simple aqueous solutions of reduced folates have overcome successfully the intact barrier of the skin models. Most impressingly the supplemented L-FTHF (L-formyltetrahydrofolate) was partially metabolized in the skin models to L-MTHF (L-methyltetrahydrofolate) the dominant form in the body and the cellular part of the skin models. The occurrence of metabolites of the added folate salts underline the accessibility for human skin cells. In addition, the scratch assay in primary keratinocytes remarkably demonstrated positive effects on wound healing. In individuals with disease-related low folate a topical folate application may help to improve skin conditions. The dermal penetration capabilities of reduced folates tested, free of any allergenic or irritating substances, open new possibilities for development of topical applications for local delivery of folate.

## References

1. Schikowski T, Hüls A. Air Pollution and Skin Aging. Curr Environ Health Rep. 2020;7:58–64. doi:10.1007/s40572-020-00262-9.

2. Parrado C, Mercado-Saenz S, Perez-Davo A, Gilaberte Y, Gonzalez S, Juarranz A. Environmental Stressors on Skin Aging. Mechanistic Insights. Front Pharmacol. 2019;10:759. doi:10.3389/fphar.2019.00759.

3. Rittié L, Fisher GJ. Natural and sun-induced aging of human skin. Cold Spring Harb Perspect Med. 2015;5:a015370. doi:10.1101/cshperspect.a015370.

4. Blair MJ, Jones JD, Woessner AE, Quinn KP. Skin Structure-Function Relationships and the Wound Healing Response to Intrinsic Aging. Adv Wound Care (New Rochelle). 2020;9:127–43. doi:10.1089/wound.2019.1021.

5. Bonté F, Girard D, Archambault J-C, Desmoulière A. Skin Changes During Ageing. Subcell Biochem. 2019;91:249–80. doi:10.1007/978-981-13-3681-2_10.

6. Pazyar N, Yaghoobi R, Rafiee E, Mehrabian A, Feily A. Skin wound healing and phytomedicine: a review. Skin Pharmacol Physiol. 2014;27:303–10. doi:10.1159/000357477.

7. Takeo M, Lee W, Ito M. Wound healing and skin regeneration. Cold Spring Harb Perspect Med. 2015;5:a023267. doi:10.1101/cshperspect.a023267.

8. Gosain A, DiPietro LA. Aging and wound healing. World J Surg. 2004;28:321–6. doi:10.1007/s00268-003-7397-6.

9. Gerstein AD, Phillips TJ, Rogers GS, Gilchrest BA. Wound healing and aging. Dermatol Clin. 1993;11:749–57.

10. Sgonc R, Gruber J. Age-related aspects of cutaneous wound healing: a mini-review. Gerontology. 2013;59:159–64. doi:10.1159/000342344.

11. Lecardonnel J, Deshayes N, Genty G, Parent N, Bernard BA, Rathman-Josserand M, Paris M. Ageing and colony-forming efficiency of human hair follicle keratinocytes. Exp Dermatol. 2013;22:604–6. doi:10.1111/exd.12204.

12. Barrandon Y, Green H. Three clonal types of keratinocyte with different capacities for multiplication. Proc. Natl. Acad. Sci. U. S. A. 1987;84:2302–6. doi:10.1073/pnas.84.8.2302.

13. Ashcroft GS, Horan MA, Ferguson MW. Aging alters the inflammatory and endothelial cell adhesion molecule profiles during human cutaneous wound healing. Lab Invest. 1998;78:47–58.

14. Ashcroft GS, Ashworth JJ. Potential role of estrogens in wound healing. Am J Clin Dermatol. 2003;4:737–43. doi:10.2165/00128071-200304110-00002.

15. Wilkinson HN, Hardman MJ. The role of estrogen in cutaneous ageing and repair. Maturitas. 2017;103:60–4. doi:10.1016/j.maturitas.2017.06.026.

16. Watson RR, Zibadi S, Preedy VR. Dietary Components and Immune Function. Totowa, NJ: Springer Science+Business Media LLC; 2010.

17. Pagano C, Perioli L, Latterini L, Nocchetti M, Ceccarini MR, Marani M, et al. Folic acid-layered double hydroxides hybrids in skin formulations: Technological, photochemical and in vitro cytotoxicity on human keratinocytes and fibroblasts. Applied Clay Science. 2019;168:382–95. doi:10.1016/j.clay.2018.12.009.

18. Hofsäss MA, Souza J de, Silva-Barcellos NM, Bellavinha KR, Abrahamsson B, Cristofoletti R, et al. Biowaiver Monographs for Immediate-Release Solid Oral Dosage Forms: Folic Acid. J Pharm Sci. 2017;106:3421–30. doi:10.1016/j.xphs.2017.08.007.

19. Pietrzik K, Bailey L, Shane B. Folic acid and L-5-methyltetrahydrofolate: comparison of clinical pharmacokinetics and pharmacodynamics. Clin. Pharmacokinet. 2010;49:535–48.

20. Kirsch SH, Knapp J-P, Herrmann W, Obeid R. Quantification of key folate forms in serum using stable-isotope dilution ultra performance liquid chromatography-tandem mass spectrometry. J Chromatogr B Analyt Technol Biomed Life Sci. 2010;878:68–75. doi:10.1016/j.jchromb.2009.11.021.

21. Schneider CA, Rasband WS, Eliceiri KW. NIH Image to ImageJ: 25 years of image analysis. Nat. Methods. 2012;9:671–5.

22. Hasoun LZ, Bailey SW, Outlaw KK, Ayling JE. Effect of serum folate status on total folate and 5-methyltetrahydrofolate in human skin. Am J Clin Nutr. 2013;98:42–8. doi:10.3945/ajcn.112.057562.

23. Organisation for Economic Co-operation and Development. Test No. 439: In Vitro Skin Irritation: Reconstructed Human Epidermis Test Method. Paris: OECD Publishing; 2015.

24. Williams JD, Jacobson EL, Kim H, Kim M, Jacobson MK. Folate in skin cancer prevention. Subcell Biochem. 2012;56:181–97. doi:10.1007/978-94-007-2199-9_10.

25. Bailey SW, Ayling JE. The extremely slow and variable activity of dihydrofolate reductase in human liver and its implications for high folic acid intake. Proc Natl Acad Sci U S A. 2009;106:15424–9. doi:10.1073/pnas.0902072106.

26. Selhub J, Rosenberg IH. Excessive folic acid intake and relation to adverse health outcome. Biochimie. 2016;126:71–8. doi:10.1016/j.biochi.2016.04.010.

27. Hiraoka M, Kagawa Y. Genetic polymorphisms and folate status. Congenit Anom (Kyoto). 2017;57:142–9. doi:10.1111/cga.12232.

28. Froese DS, Huemer M, Suormala T, Burda P, Coelho D, Guéant J-L, et al. Mutation Update and Review of Severe Methylenetetrahydrofolate Reductase Deficiency. Hum Mutat. 2016;37:427–38. doi:10.1002/humu.22970.

